# Decellularized Plant-Based Scaffolds for Guided Alignment of Myoblast Cells

**DOI:** 10.1101/2020.02.23.958686

**Authors:** Santiago Campuzano, Nicolette B. Mogilever, Andrew E. Pelling

**Affiliations:** STEM Complex, Department of Physics, University of Ottawa, Ottawa, ON, Canada; Department of Biology, University of Ottawa, Ottawa, ON, Canada; Institute for Science Society and Policy, University of Ottawa, Ottawa, ON, Canada; SymbioticA, School of Human Sciences, Physiology and Human Biology, University of Western Australia, Perth, WA 6009, Australia

## Abstract

Alignment and orientation of cells *in vivo* plays a crucial role in the functionality of tissue. A challenged faced by traditional cell culture approaches is that the majority of two-dimensional substrates fail to induce a controlled alignment of cells *in vitro*. To address this challenge, approaches utilizing mechanical stresses, exposure to electrical fields, structurally aligned biomaterials and/or textured microfabricated substrates, have been developed to control the organization of cells through microenvironmental stimuli. In the field of muscle tissue engineering it is often desirable to control the alignment and fusion of muscle precursor cells as it more closely resembles *in vivo* conditions. In this study, we utilize plant-derived cellulose biomaterials to control the *in vitro* alignment of C2C12 murine myoblasts. We hereby report that cells display a clear sensitivity to the highly aligned vascular bundle architectures found in decellularized celery (*Apium graveolens)*. Conveniently, the xylem and phloem channels lie within the 10-100μm diameter, which has been shown to be optimal diameter for myoblast alignment through contact guidance. Following 10 days in proliferation media, F-actin filaments were observed to be aligned parallel to the longitudinal axis of the vascular bundle. Subsequently, following 5 days in differentiation media, myoblast maintained an aligned morphology, which led to the formation of aligned myotubes. We therefore conclude that the microtopography of the vascular bundle guides muscle cell alignment. The results presented here highlight the potential of this plant-derived scaffold for *in vitro* studies of muscle myogenesis, where structural anisotropy is required to more closely resemble *in vivo* conditions.

## 1. Introduction

The orientation and arrangement of cells *in vivo* plays a crucial role in the functionality of tissue. ^1–3^ The multinucleated structures in muscle tissue, known as myofibers, are bundled and arranged along a predetermined axis to synchronously contract and generate force.^4,5^ In addition to skeletal muscle tissue, airways, arteries, and veins rely on the circumferential alignment of smooth muscle cells to facilitate the transport of fluids and gases.^3,6^ In the laboratory, however, *in vitro* studies are performed on flat 2D Petri dishes which lack biologically active adhesion sites, dimensionality, microtopography, and proper mechanical stimuli. This in turn causes cells to appear randomly scattered; and portray dissimilarities in proliferation, differentiation and overall gene expression.^7–13^ In order to assimilate 2D cell culture to the *in vivo* environment and further understand the role of matrix topography, substrates with a wide array of topographical structures, such as posts^14^, microchannels^15^, and nanofibers^16^ have been developed.^14^ In addition to topographical features such as microchannels/microgrooves, cyclic strain and electrical stimulation have also been shown to influence spatial orientation of cells.^17,18^

It has been shown that smooth muscle cells^11^, skeletal muscle cells^8^, neurons^10^, and tendon derived cells^12^ portrayed a difference in gene expression when compared to those cultured on the smooth surfaces of tissue culture dishes. Skeletal muscle cells upregulate troponin T, myosin heavy chain, and myogenin when cultured on uniaxial microchannels.^8^ Substrate topography has also been shown to influence differentiation lineage of mesenchymal stem cells. Mesenchymal stem cells cultured on grooves and ridges committed to myogenic and adipogenic line, whereas smooth surface induced osteogenic differentiation.^7^

Microchannel development has proven to be a popular method for guided cell alignment due to tunability and reproducibility. Photolithography^19,20^, femtosecond pulsed laser^21^, 3D printing^22,23^ and electron-beam lithography^24,25^ have all been shown to produce substrate topographies capable of inducing alignment. It has been shown that microchannel width ranging from 5-200μm can induce alignment of myoblasts, where channels 20 to 100μm wide induced optimal myotube maturation.^15,26–28^ The depth of the channels has also been shown to play a role in cell alignment. Cells cultured on 2μm deep microgrooves exhibited limited alignment after 24 h when compared to 7um deep channels, which provided a permanent alignment.^29^ This observation appeared to be cell line specific: C2C12 responded to grooves below 0.5μm differently than primary myoblasts.^27^ On the discussed substrates, cell alignment is attributed to confinement and contact guidance. Cells were considered aligned if the mean angle of cells with reference to the direction of the substrate pattern was below 10°.^27,28^

Anchorage dependent cells interact with the extracellular matrix through integrin binding, often through the RGD (arginine-glycine-aspartate) motif present in fibronectin, vibronectin and laminin.^30^ Non-animal derived scaffolds lack binding motifs; yet, the presence of adhesion proteins naturally found in serum (e.g. fetal bovine serum) and those extruded by cells facilitate interaction between a foreign matrix and the cell.^31,32^ Clusters of integrins, GTPases and other enzymes form focal adhesion (FA) complex, which undergo active restructuring in response to substrate stiffness and topography.^33,34^ Stretch sensitive channels such as Bin/amphiphysin/Rv (BAR) domain-containing proteins work in part as topographical sensors, and are believed to play a role in the sensing of curved structures.^35–37^ The reorganization of cytoskeletal proteins in response to topography is mandated in part by the regulation of Rho, Rac, and Cdc 42.^36^ This phenomenon is commonly referred to as contact guidance.^36,38–40^ Information received through focal adhesion complexes influences focal adhesion kinase (FAK) signaling, which in turn triggers downstream signaling of the mitogen activated proteins kinase (MAPK) cascade.^30,40^ Although the interaction between cells and matrix has been an area of interest for over 20 years, locomotion, local restructuring, and molecular mechanisms governing intracellular changes is still not fully understood.

A series of recent studies have depicted the biocompatibility of decellularized plant tissue *in vitro* and *in* vivo.^44,45^ Through the use of surfactants, such as SDS, the cell membrane becomes compromised leading to cell lysis.^41–44,46^ Immortalized cell lines were shown to proliferate throughout the relatively porous decellularized apple tissue without the need for biofunctionalization.^42,44^ *In vivo* studies have shown that implanted decellularized apple tissue induced a minimal immune response, and stimulated angiogenesis.^44,46^ In addition to biocompatibility, the mechanical properties have been shown to resemble that of skeletal^44^ and cardiac muscle^47^ tissue. However, decellularized plant tissue lacks the biochemical cues natively found in mammalian extracellular matrix.^48^ Yet, the lack of biochemical cues in plant cellulose, bacterial cellulose, and cellulose derivatives has not impeded applications in tissue engineering.^49,50^ Furthermore, the tunability potential of cellulose, including biofunctionalization, can further extend its applications.^41,42,44,51,52^

As part of the wide arrays of structures found in plants, the vascularization of plants is composed of vessels with diameters in the micrometer scale.^47,53–55^ In the case of celery (*Apium graveolens*), a dicot plant, the vascularization is composed of two major structures(xylem and phloem) composed of microchannels with varying diameters^53^. Therefore, based on the diameter of these structures, we hypothesized that C2C12 myoblast cells will align along the longitudinal direction of the vascular bundle. After 10 days in growth media, we found that the actin filaments and apex of nuclei align parallel to the vascular bundle. After differentiating the cells into fused myotubes, we found that myotubes maintain their align morphology.

We depict how the natural topography of the vascular bundle can be exploited to induce the uniaxial orientation of myoblast and myotubes. This presents a simple, highly accessible, biocompatible and tunable substrate for guided cell alignment. This in turn is expected to uncover new methods and applications of plant-based scaffold research and broaden our knowledge on the importance of spatial orientation and cell alignment with respect to unconventional biomaterials. Although we only discuss skeletal muscle cells, spatial alignment and orientation is an important property found in all developing and mature tissues. ^1–3,56^ The work presented here points to the potential use of the vascular bundles in plant tissues as a simple tool for further development of microenvironments for alignment and orientation studies of other cell types.

## 2. Materials and Methods

### 2.1 Scaffold Preparation

The decellularization protocol was performed as described previously.^44^ Briefly, celery (*Apium graveolens)* stalk was cut parallel and perpendicular to the longitudinal axis using a mandolin slicer. A 6mm biopsy punch was then used to obtain round scaffolds with the exposed vascular bundle in a longitudinal and cross section orientation. (Fig.1A). The samples were then transferred to a 15mL Falcon tube containing 0.1% SDS at a ratio of one sample per mL. Samples were then agitated in a shaker at 120RPM for 72 hours. Following treatment with SDS, the samples were washed three times with deionized water. After the final wash, 100mM solution of CaCl_2_ (1mL/scaffold) was added and samples were incubated at room temperature for 24 hours. After 24 hours, the samples were washed with distilled water three times. On the final wash, the water was removed and 70% ethanol was added for 30min. At this point, the samples were brought into a class II biosafety cabinet and washed three times with sterile PBS. The samples were placed on PDMS coated 12-well plates with 2mL of growth media (refer to next section). The samples were incubated overnight at 37°C and 5% CO_2_. Prior to cell seeding, the media was removed (Fig.1A).

**Figure 1.**
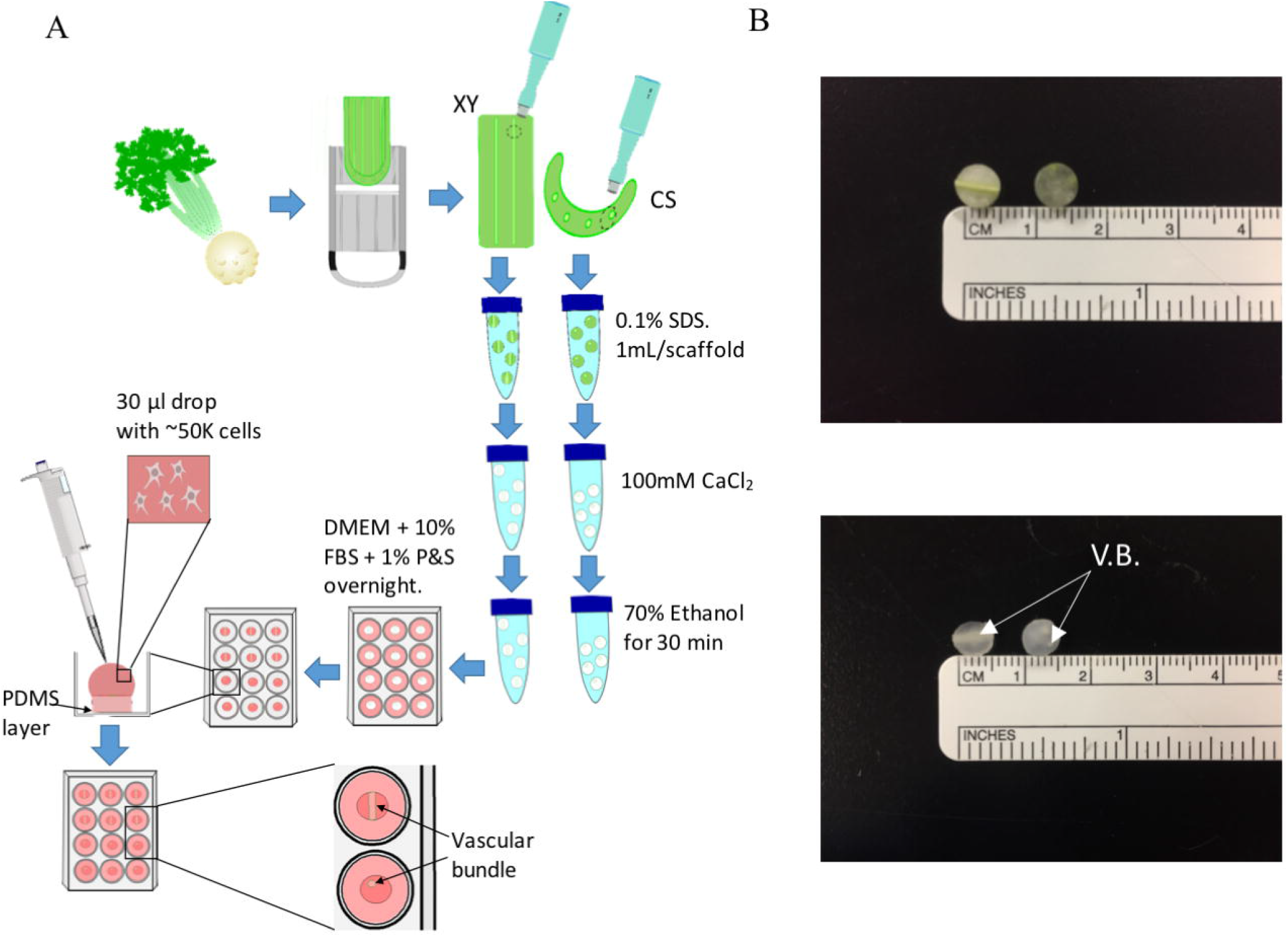
(A) Visual representation of celery(*A. graveolens)* scaffold preparation. (B) Samples were 6mm wide and 2.15±0.15mm thick. “XY” corresponds to scaffolds cut (left) longitudinally with respect to the celery stalk, whereas” CS” corresponds to (right) cross sections. Samples became clear following the 3-day incubation period in 0.1% SDS. Approximately 50,000 cells were seeded on (C) decellularized scaffolds and left on scaffold for 4.5 hours. V.B.= Vascular bundle.

### 2.2 Cell culture

C2C12 murine myoblasts were plated on tissue culture plates and maintained at 37°C with 5% CO_2_. Cells were cultured in growth media consisting of high glucose DMEM with L-glutamine and sodium pyruvate (Hyclone) supplemented with 10% FBS (Wisent) and 1% penicilin (10000U/mL) & streptomycin (10000ug/mL) (Hyclone). Once the cells reached 70-80% confluency, they were trypsinized (0.05%), resuspended in growth media, and spun down in the centrifuge at 1000RPM (97g) for 3 min. Following centrifugation, the pellet was resuspended in growth media to acquire 1.7 × 10^6^ cells/mL. Cells were counted using a hemocytometer and trypan blue to determine viability. 30 μL of media containing cells were placed on the scaffolds and incubated for 4.5 hours. Following incubation, 2mL of growth media was added and samples were incubated for the 10 days with media change every 48 hours until day 7, followed by daily media change until day 10. For differentiation studies, cells were placed in differentiation media for 5 days. The differentiation media was composed of high glucose DMEM, 2% horse serum (Gibco) and 1% penicillin and streptomycin (Hyclone).

### 2.3 Fluorescent staining

Following 10 days in growth media, decellularized scaffolds with cells were transferred to a microcentrifuge tube using a metal paddle (minimize contact with exposed vascular bundle) and washed three times with PBS. The samples were fixed with 3.5% paraformaldahyde in 2% aqueous sucrose solution for 10min and washed three times with PBS. Following the final PBS wash, room temperature triton-X100 was added to permeabilize the cells. The scaffolds were once again washed three times with PBS. For F-actin imaging, scaffolds were stained using Alexa Fluor 488 phalloidin (Invitrogen) in PBS at a 1:200 concentration and incubated for 20min in the dark. Nuclei was stained by first placing the scaffolds in 10% RNAse in PBS (DNase and protease-free) (Thermo Fisher) for 30 min at 37°C, followed by PBS wash (3X). After the third wash, propidium iodide(1mg/mL) (Thermo Fisher) was added at a 1:1000 concentration for 30min. In order to observe the cells morphology with respect to the substrate, cellulose was stained with 10% calcofluor in PBS for 20min at room temperature. Prior to seeding, native and decellularized scaffolds were placed in 1:500 Hoescht 33342(Invitrogen) made up in phosphate buffered saline (PBS) for 30min at 37°C to confirm successful decellularization.

To test for the presence of myotubes, scaffolds from the 5-day differentiation treatment were fixed and permeabilized as described above. These samples were then washed three times with cold wash buffer (5% FBS in PBS) and placed at 4°C for 20min. The cold wash buffer was removed and MF-20 (DSHB Hybridoma Product) was added at a 1:200 concentration made up in cold wash buffer and incubated for 24 hours at 4°C. The MF-20 solution was removed and the scaffolds were washed three times with cold wash buffer. Samples were stored in cold wash buffer for 20 min at 4°C before adding the secondary antibody. Anti-Mouse IgG (whole molecule)–FITC (Sigma) was added at a 1:100 concentration and placed in the refrigerator for 1 hour. The samples were then washed 3 times with PBS prior to imaging.

### 2.4 Microscopy

Scaffolds were placed on coverslips with mounting medium (Vectashield H-1000) prior to imaging. The samples were imaged with a Nikon TiE A1-R high speed resonant scanner confocal microscope with a 10X and 40× lens. Image processing was done on FIJI- ImageJ. The images presented throughout this manuscript are maximum projections of 210 ± 51μm (N=9) and 49 ± 24μm (N = 7) confocal volumes for 10X and 40X magnification, respectively. Brightness of fluorophore signal was enhanced to improve contrast of structures.

### 2.5 Scanning electron microscopy

Sample preparation was performed as described previously.^57^ Briefly, the sample was washed three times with PBS and placed in aqueous solution containing 3.5% paraformaldehyde (in 2% sucrose solution) and 1.5% glutaraldehyde (Final concentration) overnight. Post fixation, the sample was washed again with PBS and dehydrated through a sequential ethanol gradient (30,50,75,95 & 99%). The sample was dried using the Samdri-PVT-3D Critical point dryer in 99% ethanol. The dried sample was gold sputtered with a 5nm layer (LEICA EM ACE 200). The samples were image with a JEOL JSM-7500F FESEM at 2.0 KV.

### 2.6 Orientation measurement

1.61mm^2^ images with a 24 ± 6% (N=20) actin coverage and 20 ± 10% (N=20) MYHC coverage were analyzed to determine the orientation of the labeled structures. The images were captured 1mm away from the edge of the vascular bundle to minimize the influence of damaged channels brought upon by the biopsy punch and mandolin. The labeled MYHC and actin were first thresholded with the *Adaptive Threshold* plug-in on ImageJ-FIJI to isolate labeled cytoskeletal proteins and vascular bundle. The directionality of the structures was determined using the *Directionality* plug-in on FIJI ImageJ. We quantified the degree of alignment by examining the broadness of the Gaussian distribution. In a manner similar to signal processing techniques, we defined the alignment in a manner reminiscent of a Quality Factor(Q-factor). In this case, the Degree of Alignment = Average Angle / Width of the distribution at half maximum. Tightly aligned myotubes will therefore have a small distribution and large degree of alignment concomitantly with a large Degree of Alignment. Conversely, a population of myotubes which are randomly oriented will have a broad distribution and a small Degree of Alignment.

### 2.7 Histology

The scaffolds were rinsed with PBS three times and fixed overnight with 10% formalin (Sigma). Following formalin fixation, the scaffolds were rinsed with PBS and stored in 70% ethanol until paraffin embedding. The vascular bundle was serial cut into 4μm sections (longitudinally) and stained with H&E. The slides were scanned using the Aperio AT2 pathology slide scanner. Image viewing and processing was done with Aperio ImageScope.

### 2.8 Immunohistochemical analysis

Formalin-fixed, paraffin-embedded sections were deparaffinized in xylene and rehydrated through a sequential 100-70% ethanol gradient. Heat induced epitope retrieval was done at 110°C for 12 min with citrate buffer pH 6.0. Sections were then blocked with Rodent Block M blocking reagent (BioCare) and incubated overnight at 4°C at a 1:12 dilution using MF-20 antibody (DSHB). Following overnight incubation, sections were washed with 1xTBST and then incubated with goat anti-mouse-568(Abcam) secondary antibody at a 1:500 dilution for 2 hours in the dark at room temperature. Sections were washed with a 1X TBST, incubated for 5 min with a quencher (Vector TrueView Autofluorescence Quenching, Vector Labs). Sections were washed and incubated with 5ug/mL DAPI (ThermoScientific), washed and cover slipped.

### 2.9 Migration Measurement and channel diameter

In order to determine the migration of cells throughout the vascular bundle, nuclei on longitudinal cuts 12 μm apart were counted to minimize the probability of counting the same nuclei twice or underestimating the actual number. This was performed based on an average nuclei size of 10 μm.^58^ An Open Computer Vision (OpenCV) Python script was developed to batch process images. The location of each nuclei was then determined by exporting the coordinates of a mouse-click. Once a nucleus was counted, an overlay green dot would mark it to prevent overestimation (https://github.com/pellinglab/Vessel-Diameter). Similarly, a separate OpenCV Python script was also used to measure the diameter of the vascular bundle channels. The script first isolates the edges and then draws built-in Contours. The contour area was fitted to a circle based on the correlation between diameter and surface area (https://github.com/pellinglab/Vessel-Diameter).

### 2.10 Statistical analysis

Student’s t-test assuming equal variance was used to analyze the results presented in this manuscript. All of the statistical tests were done on R studio. Alpha value was set at 0.05. Values throughout this manuscript are displayed as mean ± standard deviation.

## 3. Results

### 3.1 Substrate preparation

We first sought to create two distinct scaffolds with different topographies. The scaffolds were created by cutting the *A. graveolens* stalk longitudinally and cross-sectionally (Fig.1A). Cross sections had an amorphous topography, whereas the longitudinal scaffolds portrayed highly aligned vascular bundle grooves. Following 3 days in 0.1% SDS, the scaffolds lost their green color due to the loss of cellular components (Fig.1B). In order to facilitate removal of SDS, 100mM concentration of CaCl_2_ was added to the scaffolds for 24 hours. The decellularization of the vascular bundle was determined using a membrane permeable stain, Hoescht 33342 (Fig.S1). A comparison between native and decellularized phloem was expected to depict the nuclei of companion cells in the phloem, as opposed to the outline of companion cells lacking a nuclei in the decellularized scaffolds. The xylem and sieve tube elements don’t possess a nucleus.^59^ The loss of the natural green colour and nuclei of companion cells depict successful decellularization of the *A. graveolens* scaffolds.

The lignified tissue in the xylem interacts with propidium iodide leading to the observed red fluorescence. In contrast, the parenchyma and phloem labelled with calcofluor led to blue fluorescence (Depicted in Fig.2A & Fig.2B as yellow and purple, respectively). The diameter of the phloem vessels was determined to be 16 ± 6μm (N=3), whereas the xylem vessels were 17 ± 5μm (N=3) wide. Based on these results, we conclude that were able to acquire grooves with diameters within the 10-100μm region necessary for optimal alignment.

**Figure 2.**
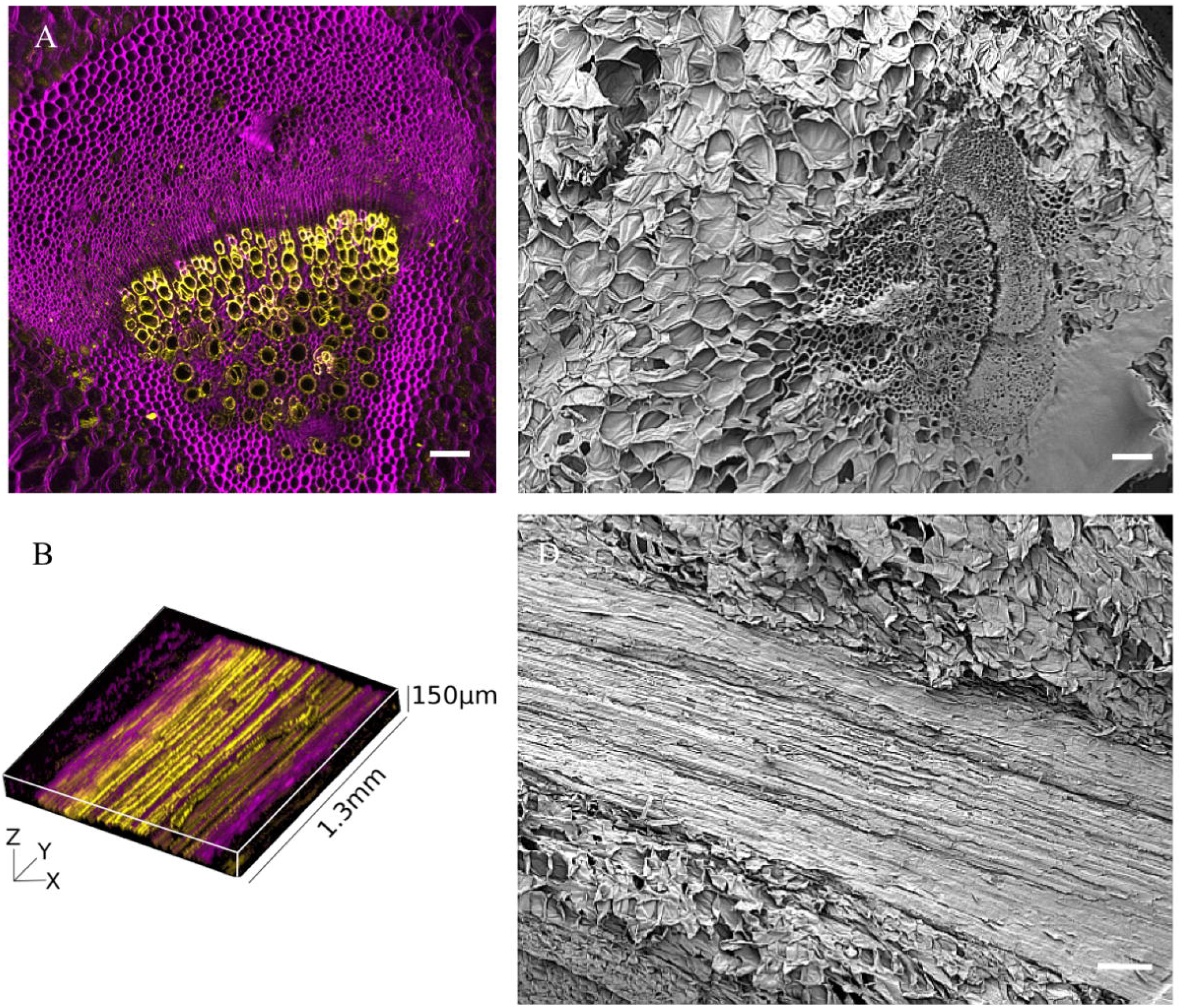
Vascular bundle of *A. graveolens*. The vasculature of celery is composed of channels in the micrometer scale, which are segregated into two major structures: phloem(16 ± 6μm. N=3) and xylem (17 ± 5μm. N=3). A) Max projection of cross section and (B) 3D reconstruction of longitudinally cut vascular bundle. Phloem and ground tissue(parenchyma) were labeled with calcofluor(purple); lignified tissue was stained with propidium iodide(yellow). SEM images of vascular bundle (C) cross section and (D) longitudinal cut. (A) Scale bar = 100 μm. (C&D) Scale bar = 200 μm.

### 3.2 F-actin alignment on vascular bundle derived from *A. graveolens* stalks

We next sought to test our hypothesis that microscale grooves in a plant scaffold would lead to cellular alignment. Cells were seeded on the decellularized scaffolds at a concentration of ~5 × 10^5^ cells/mL. A 30μl drop of cells was left on the scaffold for 4.5 hours. The scaffolds were incubated in growth media for 10 days. By the end of the incubation period, F-actin filaments were observed to be oriented parallel to the longitudinal axis of the vascular bundle (Fig.3F,3J) as opposed to those on the parenchyma (Fig.3B). This in turn leads to a visible unimodal distribution (Fig.3M) (0.5 ± 41.2) of aligned F-actin filaments, whereas F- actin filaments on the parenchyma are represented by a uniform histogram (Fig.3N) (0.7 ± 50.2°). In this case, the histograms represent either a predominant orientation angle (anisotropy) or uniform distribution of F-actin filaments(isotropy). Furthermore, a significant difference was observed between the Degree of Alignment of myoblast seeded on the vascular bundle and those on the parenchyma. (Aligned myoblast Degree of Alignment: 4.7 ± 2.7. Isotropic myoblast Degree of Alignment: 0.7 ± 0.4. P = 2 × 10^−4^, N = 10). Consequently, the normalized mean value of F-actin filaments with respect to the vascular bundle’s longitudinal axis was determined to be 1.2° ± 2.0 (N =10). This led us to conclude that the vascular bundle topography had a clear influence in the phenotype and reorganization of F-actin filaments.

**Figure 3.**
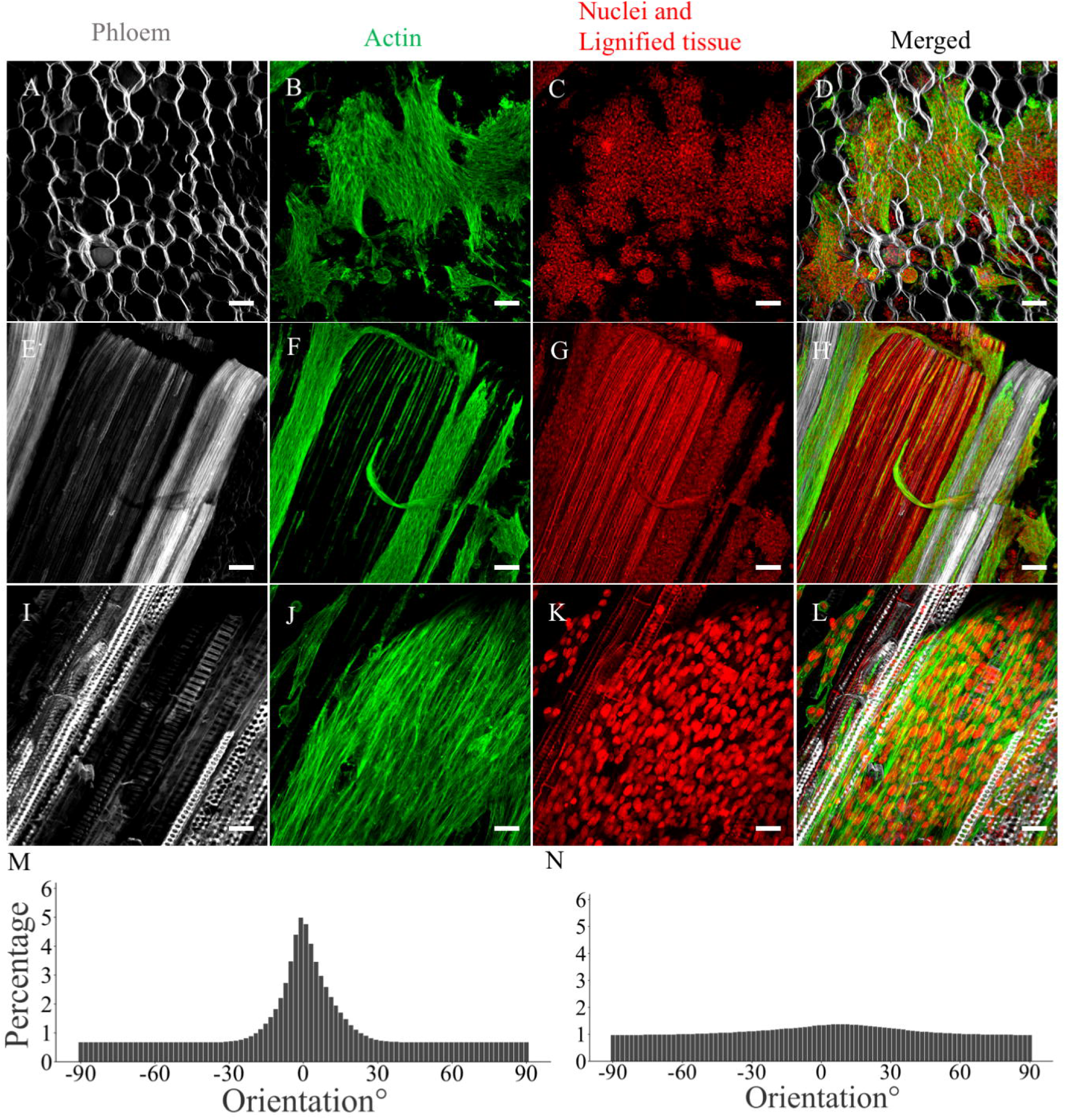
Myoblast alignment on the decellularized vascular bundle of celery (*A. graveolens)* and celery parenchyma following 10 days in growth media. As opposed to the amorphous topography of plant parenchyma, the vascular bundle provides a highly aligned microtopography capable of inducing myoblast alignment hereby depicted by the orientation of F-actin filaments. (A,E & I) Phloem and ground tissue were labeled with Calcoflour (gray); (B, F, & J) F-actin was labeled with Phalloidin Alexa fluor 488(green) ; (C, G & L) nuclei and lignified tissue were labeled with propidium iodide (Red) (A - H) Scale bar = 100μm. (I-L) Scale bar = 25μm. Histograms representing average F-actin filament orientation on (M) vascular bundle and (N) parenchyma (N=10 max projection images).

### 3.3 Fusion of aligned myoblast on vascular bundle derived from *A. graveolens*

The clear reorganization of F- actin filaments and the orientation of the nuclei-apex led us to hypothesize that 5 days in differentiation will lead to the formation of aligned myotubes. Cells were first incubated for 10 days in proliferation media, and then switched to differentiation media for 5 days to induce fusion; yet, minimize the risk of detachment induced by active contractions.^60–63^. Myosin heavy chain (MYHC), an early differentiation marker, was labeled with the MF-20 antibody (Fig.4B,4F,4J). The average myotube length was determined to be 308 ± 169μm (N=103). Myotubes were predominantly observed to have an elongated morphology (Fig.4F,4J). Concomitantly with the F-actin orientation, a statistically significant difference was also observed between the orientation of the myotubes cultured on the vascular bundle (Fig.4F,4J) and those on the parenchyma (Fig.4B) (aligned myotubes Degree of Alignment: 3.3 ± 1.1. Isotropic myotubes Degree of Alignment: 0.6 ± 0.6. P = 2 × 10^−6^, N =10). Concomitantly with F-actin filament orientation, a unimodal histogram with a broader distribution of 1.7 ± 45.5° represents the orientation of myotubes on the vascular bundle with reference to the long-axis of the vascular bundle (Fig.4M). Conversely, and in support of isotropic F-actin filament formation on celery parenchyma, a uniform histogram (−4.4 ± 51.1°) depicts the isotropic formation of myotubes on celery parenchyma (Fig.4N). In addition, the normalized mean value for the orientation of myotubes with respect to the vascular bundle was calculated to be 8.6° ± 23.8 (N = 10). Taking into consideration the natural variation of biological samples and the preparation method, we can’t disregard the formation of smooth or damaged areas with outlying topographical cues or void thereof (Fig.S2).

**Figure 4.**
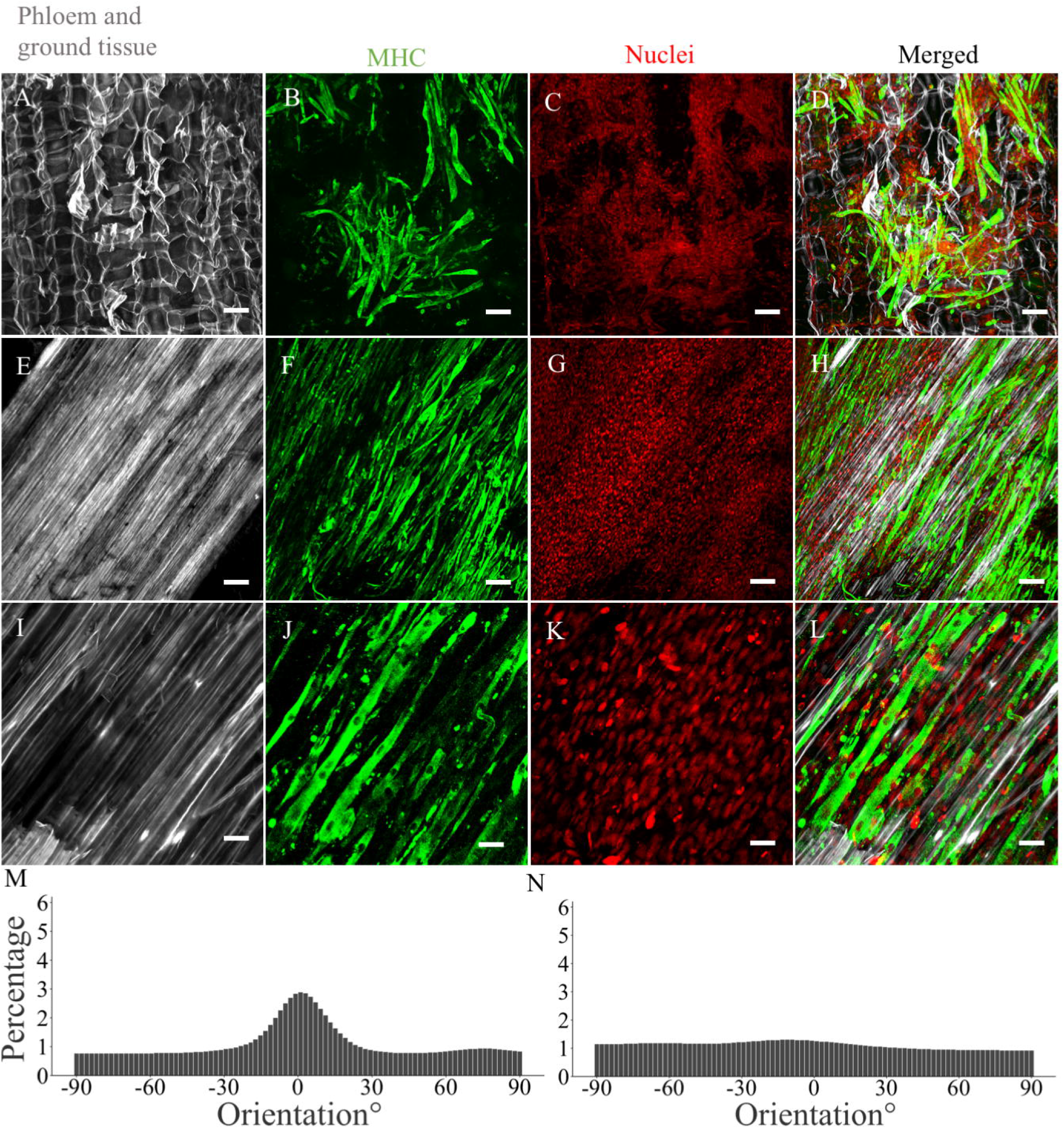
Myotube alignment on the decellularized vascular bundle of celery(*A. graveolens)* and celery parenchyma. Myoblast maintain an aligned morphology through the fusion (myogenesis) process, leading to the formation of aligned myotubes. Myotubes were labeled with MF-20 followed by FITC-conjugated secondary antibody (green); nuclei were labeled with propidium iodide (Red); ground tissue and vascular bundle were labeled with calcofluor (gray). Scale bar = 100μm. Histograms representing average myotube orientation on (M) vascular bundle and (N) parenchyma(N = 10 max projection images).

### 3.4 Migration of C2C12 through vascular bundle and immunohistochemical analysis of MYHC expression

In addition to studying the effects of alignment, we lastly sought to determine if cells seeded on cross section scaffolds would be able to migrate the full extent of the vascular bundle. Therefore, cells were seeded on the decellularized scaffolds as mentioned previously and incubated for 10 days. The scaffolds were cut cross sectionally (Fig.5B,5E) and longitudinally (Fig.5A,5F). The average migration distance was determined to be 0.91 ± 0.80mm (N=5)(Fig.5C). Cells were predominantly observed on the surface of the vascular bundle as depicted by the bars at the 0 and 2mm mark. These surface-cells correspond to those that didn’t migrate or migrated fully across the vascular bundle, respectively. Throughout the bundle, however, cells appear to be uniformly distributed (Fig.5C). Cells present an elongated phenotype (Fig.5A,5D), which corresponds to optimal cell-matrix interaction. Following the observed distribution of cells throughout the vascular bundle after 10 days in growth media, the cross-section scaffolds were placed in differentiation media for 5 days to induce myoblast fusion. Based on immunohistochemical analysis of MYHC expression, myotube formation was not observed (Fig.S3).

**Figure 5.**
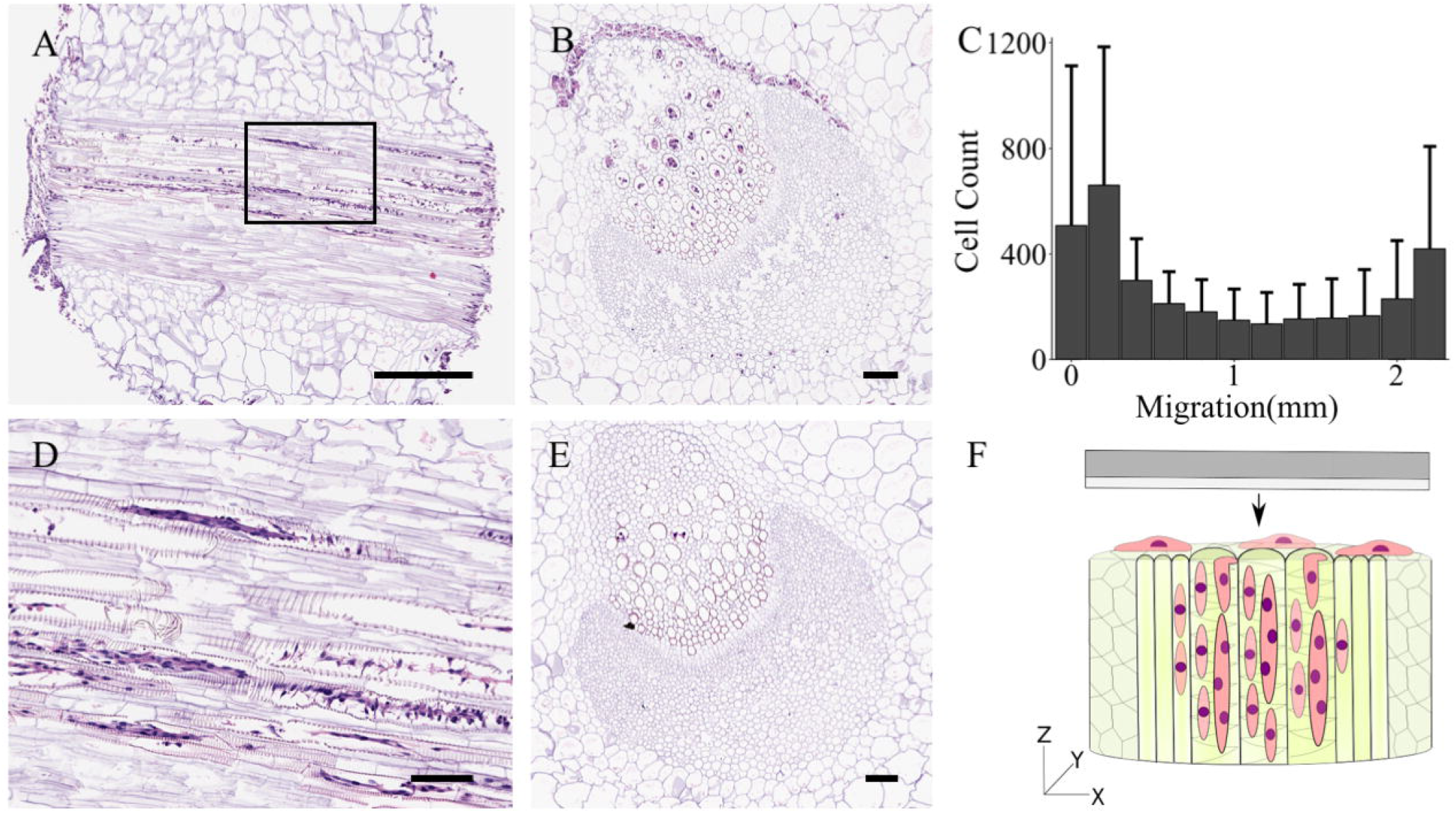
Migration of C2C12 through decellularized vascular bundler of celery at day 10. A) 4μm wide longitudinal cut of vascular bundle. (F) visual representation of longitudinal cut. B) Cross section of vascular bundle 100um away from surface. (E) cross section of vascular bundle 800um away from surface. Cells were stained with H&E. C) Migration distribution of cells throughout the vascular bundle after a 10-day incubation period (N=5)(Mean ± S.D). A) Scale bar = 500μm. (B-D) Scale bar = 100μm.

## 4. Discussion

Alignment and spatial orientation of cells *in vivo* correlates strongly with tissue functionality.^1–3^ In the case of skeletal muscle tissue, muscle precursor cells proliferate, fuse along a predetermined axis, and form fascicles composed of striated myofibers.^4^ Depolarization of these myofibers leads to the contractions that drive motion in birds^64^, amphibians^65^, crustaceans^66^, and mammals.^67^

In order to better understand skeletal muscle tissue, skeletal muscle cells are routinely cultured in stiff and smooth 2D Petri dishes. By underrepresenting the *in vivo* environment, cells appear randomly scattered and display a significant difference in gene expression.^4,68^ In order to recreate alignment *in vitro*, fibers^8,69,70^, microchannels of various sizes^15,20,71^, cyclic strain^17,72^, and electrical stimulation^18^ have been use to influence spatial orientation. *In-vitro* studies have showed that myoblast alignment upregulated the expression of troponin T, myogenin, and myosin heavy chain II.^8^ Another group showed that cell viability and proliferation of smooth muscle cells increased on aligned PHBV nanofibers, alongside an increase in gene expression of contractile markers.^11^

Great emphasis has been placed on microchannel development techniques, such as 3D printing^22^, electron-beam lithography^25^, photolithography^15,20,29^, and softlitography^73^ due to reproducibility and tunability. Yet, microchannel development is a laborious endeavor which often relies on specialized equipment. Studies on microchannel-influenced cell alignment have shown that 5-200um wide channels significantly influence the spatial orientation of skeletal muscle cells, where 10-100μm wide channels produce optimal alignment and myotube formation.^15,26^. The vascularization of plants presents a promising topography for guided cell alignment.^41^ In the case of celery (*A. graveolens)*, the vascular bundle is composed of xylem (which includes cambium) and phloem. The xylem is composed of 17 ± 5μm (N=3) wide channels, whereas the phloem is composed of 16 ± 6μm (N=3) wide channels (Fig.2A). By cutting the *A. graveolens* stalk longitudinally, we were able to acquire grooves with diameters within that of intact channels. The decellularization of the *A. graveolens* scaffold was done based on previously published protocol (Fig.1A).^44^ As show previously by a number of groups, decellularized plant tissue can be used as a substrate for 3D cell culture of immortalized and primary mammalian cells.^41,42,44,47^.

Following 10 days in culture, the actin filaments and nuclei (direction of apex) of C2C12s were observed to be oriented parallel to the longitudinal axis of the V.B when compared to cells on seeded on the parenchyma (P = 2 × 10^−4^, N = 10) (Fig.3). Taking into consideration the non-uniform arrangement of channels within the vascular bundle leads us to assume that the diameter of the channels varied. It’s then safe to say that in some cases cells were confined and in other cases guided through contact guidance. As reported by Altomare et al. (2010), 25 μm and 50μm wide grooves with a depth between 0.5 and 2.5μm presented enough of a topographical cue for cell alignment. This observation, however, appeared to be cell line specific: C2C12 didn’t respond to grooves below 0.5μm as well as primary myoblast.^27^

Consideration of myoblast alignment led us to hypothesize that myotubes would also form parallel to the longitudinal axis of the V.B. Following 5 days in differentiation media, the direction of analyzed myotubes was observed to be significantly influenced by the substrate (P = 2 × 10^−6^, N =10) (Fig.4). Myotubes on ground tissue yielded a mostly uniform histogram, whereas myotubes with an average length of 308 ± 169μm (N=103) seeded on the V.B. yielded a skewed histogram. The images analyzed here yielded a mean normalized direction of 1.2 ± 2.0° and 8.6 ± 23.8° (N = 10) for F-actin and myotubes, respectively. With reference to literature values, cells were considered to be aligned when the normalized direction of myotubes with respect to the substrate was below 10°.^27,28^ This further supports our observation that the microtopography of the vascular bundle induced guided alignment of C2C12 murine myoblast. The spread of the myotube-histogram is likely due to the noise from non-specific antibody binding, lack of topographical cues (Fig.S2) and partial detachment of cells during the staining steps.

In addition to alignment, we aimed to determine if cells would migrate across a 2.15±0.15mm long V.B. Histological analysis shows a uniform distribution of cell throughout the length of the scaffold with a greater number of cells on the surface (Fig.5C). Although cells were uniformly distributed throughout the bundle (Fig.5C), myotube formation was not observed following 5 days in differentiation media (Fig.S3). Confluency is a crucial factor in myogenesis; therefore, to obtain optimal myotube development *in vitro*, differentiation media should be introduced into an 85-95% confluent myoblast culture.^74^ Consequently, MYHC expression has been shown to be influenced by cell density and confluency.^75^ We therefore conclude based on the samples tested that confluency was suboptimal for myotube formation. We recommend future studies into myogenesis on vascular bundles to introduce longer proliferation periods.

Here we show that the vascular bundle of *A. graveolens* is able to induce alignment of myoblast and subsequently myotubes. However, it lacks a wide array of factors that influence cells *in vitro*, such as biochemical and mechanical cues. As elucidated previously, the xylem and phloem of plants has been determined to be approximately 10^6^ times^76,77^ and 10^3^ times^78^ stiffer than muscle tissue^79^, respectively. With reference to mammalian cells, stiffness has been shown to influence cell behavior, such as viability, morphology and differentiation.^80,81^ This in turn explains the outlying occurrence (Fig.4D), which portrays a lower number of cells on the xylem compared to the phloem. In addition to the 1000-fold difference in stiffness, we can’t disregard the natural hydrophobicity of lignin.^82^ As reported by Papenburg and colleagues, hydrophobic surfaces have been shown to improve initial cell attachment of C2C12 pre-myoblast; yet, a reduced proliferation rate and spreading was observed.^83^

In this study, the expression of MYHC was tested with a non-isoform specific antibody, MF-20. The antibody presented here recognizes all MHC isoforms. Therefore, this doesn’t provide an insight into the differentiation stage of myotubes. Without testing for other markers, such as Acta1 and Glut4, its difficult to speculate on the stage of myogenesis.^4^ Subsequently, based on the centralized location of the nuclei, as opposed to that of mature muscle tissue where the nuclei are found in the periphery^57,84^, it’s safe to say that the myotubes were still immature. It’s still unclear if C2C12 cultured on the decellularized vascular bundle of *A. graveolens* can express sarcomeric proteins. Yet, we can’t however, disregard the notion that striated myotubes are rarely observed *in vitro*, especially in immortalized cell lines.^4,61,62^

Microchannel fabrication often involves post treatment and coating with bioactive factors to increase cell adhesion.^20,24,71,85^ In contrast, the vascular bundle allowed for adherence and differentiation of muscle cells without a need for a bioactive coating. Yet, we must take into account the role of adhesive proteins naturally found in FBS.^31,86^ We do however postulate that biofunctionalization will improve the substrates biocompatibility and extend the use of this abundant, simple and biocompatible material to more problematic cell types.^41^ Lastly, we speculate that cells seeded on top of already aligned myotubes can lead to 3D tissue development as shown previously.^15^

As mentioned previously, the need for anisotropy in 2D cell culture doesn’t only apply to myotubes, but also neurons^10^, smooth muscle cells^11^, and tendon derived cells^12^. In addition to somatic cells, the topography presented here can be used to further study the influence of topographical cues on stem cell differentiation.^7^ We hereby present a method that will facilitate *in vitro* research on the importance of cellular anisotropy and spatial orientation.

## 5. Conclusion

Here we present a simple, reproducible, abundant, and non-animal derived substrate for guided alignment of C2C12 murine myoblast. By longitudinally cutting the vascular bundle of decellularized celery (*A. graveolens)*, we were able to acquire grooves in the micrometer scale. After culturing cells in these substrate for 10 days, F-actin filaments and apex of nuclei were observed to be aligned in the direction of the grooves. Subsequently, 5 days in differentiation media led to myoblast fusion alongside MHC expression. Yet, it’s still unclear whether myoblast cultured on this substrate have the potential to develop a functional sarcomere.

## Acknowledgement

The Authors thank Dr. Yun Liu for the assistance with SEM and Mrs Zaida Ticas for assistance with Histology. This work was supported by the New Harvest grant number 0005 and the In Vivo Foundation. New Harvest is a 501c3 non-profit research institute based in the US. S.C was also supported by the Queen Elizabeth II graduate scholarships in science and technology (QEII - GSST). We also acknowledge generous support from the Natural Sciences and Engineering Research Council (NSERC) Discovery Grant and Canada Foundation for Innovation grant.

## References

1. Y. Feng, R. J. Okamoto, R. Namani, G. M. Genin and P. V. Bayly, J. Mech. Behav. Biomed. Mater., 2013, 23, 117–132.

2. W. R. Frontera and J. Ochala, Calcif. Tissue Int., 2015, 96(3), 183–195.

3. T. Komuro, J. Desaki and Y. Uehara, Cell Tissue Res., 1982, 227(2), 429–437.

4. J. Chal and O. Pourquié, Development, 2017, 144(12), 2104–2122.

5. T. Narayanan, V. Lombardi, L. Lucii, M. Reconditi, Y. Sun, M. Linari, … M. Irving, Nature, 2002, 415(6872), 659–662.

6. J. F. Clark and G. Pyne-Geithman, Pathophysiology, 2005, 12(1), 35–45.

7. P. Wang, W. Li, J. Yu and W. Tsai, J. Mater. Sci.: Mater. Med., 2012, 23(12), 3015–3028.

8. A. Cooper, S. Jana, N. Bhattarai and M. Zhang, J. Mater. Chem., 2010, 20(40), 8904–8911.

9. P. Mozetic, S. M. Giannitelli, M. Gori, M. Trombetta and A. Rainer, J. Biomed. Mater. Res., Part A, 2017, 105(9), 2582–2588.

10. J. M. V. Basso, M. Simon and C. Staii, MRS Commun., 2018, 8(2), 487–492.

11. P. Kuppan, S. Sethuraman and U. M. Krishnan, Int. J. Polym. Mater. Polym. Biomater, 2016, 65(16), 816–825.

12. J. Foolen, S. L. Wunderli, S. Loerakker, J. G. Snedeker, Matrix Biol., 2018, 65, 14–29.

13. H. Gao, X. Cao, H. Dong, X. Fu and Y. Wang. J. Mater. Sci. Technol. (Shenyang, China), 2016, 32(9), 901–908.

14. N. Goedecke, M. Bollhalder, R. Bernet, U. Silvan and J. Snedeker, J. Visualized Exp., 2015, 105, e53350.

15. S. L. Hume, S. M. Hoyt, J. S. Walker, B. V. Sridhar, J. F. Ashley, C. N. Bowman and S. J. Bryant, Acta Biomater., 2012, 8(6), 2193–2202.

16. T. Fee, S. Surianarayanan, C. Downs, Y. Zhou and J. Berry, PloS One, 2016, 11(5), e0154806.

17. B. Liu, M. Qu, K. Qin, Z. Li, H. Li, B. Shen and Z. Jiang, Biophys. J., 2008, 94(4), 1497–1507.

18. T. Tanaka, N. Hattori-Aramaki, A. Sunohara, K. Okabe, Y. Sakamoto, H. Ochiai, … K. Kishi, International Journal of Tissue Engineering, 2014, DOI:10.1155/2014/621529.

19. P. Camelliti, P. Kohl, A. D. McCulloch and J. O. Gallagher, Nat. Protoc., 2006, 1(3), 1379–1391.

20. A. Leclerc, D. Tremblay, S. Hadjiantoniou, N. V. Bukoreshtliev, J. L. Rogowski, M. Godin and A. E. Pelling, Biomaterials, 2013, 34(33), 8097–8104.

21. W. Y. Yeong, H. Yu, K. P. Lim, K. L. G. Ng, Y. C. F. Boey, V. S. Subbu and L. P. Tan, Tissue Eng., Part C, 2010, 16(5), 1011–1021.

22. Z. Tan, T. Liu, J. Zhong, Y. Yang and W. Tan, J. Biomed. Mater. Res., Part A, 2017, 105(12), 3281–3292.

23. A. Tijore, S. A. Irvine, U. Sarig, P. Mhaisalkar, V. Baisane and S. Venkatraman, Biofabrication, 2018, 10(2), 025003.

24. P. Wang, H. Yu and W. Tsai, Biotechnol. Bioeng., 2010, 106(2), 285–294.

25. N. Idota, T. Tsukahara, K. Sato, T. Okano and T. Kitamori, Biomaterials, 2009, 30(11), 2095–2101.

26. Y. Sun, R. Duffy, A. Lee and A. W. Feinberg, Acta Biomater., 2013, 9(8), 7885–7894.

27. L. Altomare, N. Gadegaard, L. Visai, M. C. Tanzi and S. Farè, Acta Biomater., 2010, 6(6), 1948–1957.

28. J. L. Charest, A. J. García and W. P. King, Biomaterials, 2007, 28(13), 2202–2210.

29. Y. Zhao, H. Zeng, J. Nam and S. Agarwal, Biotechnol. Bioeng., 2009, 102(2), 624–631.

30. I. A. Janson and A. J. Putnam, J. Biomed. Mater. Res., Part A, 2015, 103(3), 1246–1258.

31. E. G. Hayman, M. D. Pierschbacher, S. Suzukia and E. Ruoslahti, Exp. Cell Res., 1985, 160(2), 245–258.

32. L. V. Turoverova, M. G. Khotin, N. M. Yudintseva, K. Magnusson, M. I. Blinova, G. P. Pinaev and D. Tentler, Cell and Tissue Biology, 2009, 3(5), 497–502.

33. M. Wozniak, Biochim. Biophys. Acta, Mol. Cell Res., 2004, DOI:10.1016/S0167-4889(04)00099-0.

34. C. Wu, Cell Adhes. Migr., 2007, 1(1), 13–18.

35. C. Mim and V. M. Unger, Trends Biochem. Sci., 2012, 37(12), 526–533.

36. K. Kulangara and K. W. Leong, Soft Matter, 2009, 5(21), 4072–4076.

37. M. Sheetz and V. Vogel, Nat. Rev. Mol. Cell Biol., 2006, 7(4), 265–275.

38. P. Linke, R. Suzuki, A. Yamamoto, M. Nakahata, M. Kengaku, T. Fujiwara, M. Tanaka, Langmuir, 2019, 35(23), 7538–7551.

39. A. I. Teixeira, G. A. Abrams, P. J. Bertics, C. J. Murphy and P. F. Nealey, J. Cell Sci., 2003, 116(Pt 10), 1881–1892.

40. D. Kim, P. P. Provenzano, C. L. Smith and A. Levchenko, J. Cell Biol. 2012, 197(3), 351–360.

41. G. Fontana, J. Gershlak, M. Adamski, J. Lee, S. Matsumoto, H. D. Le, … W. L. Murphy, Adv. Healthcare Mater., 2017, 6(8), 1601225.

42. D. J. Modulevsky, C. Lefebvre, K. Haase, Z. Al-Rekabi and A. E. Pelling, PLoS One, 2014, 9(5), e97835.

43. R. B. Brown and J. Audet, J. R. Soc., Interface, 2008, 5(Suppl 2), S131–S138.

44. R. J. Hickey, D. J. Modulevsky, C. M. Cuerrier and A. E. Pelling, ACS Biomater. Sci. Eng., 2018, 4(11), 3726–3736.

45. S. Campuzano and A. E. Pelling, Frontiers in Sustainable Food Systems, 2019, 3, 38.

46. D. J. Modulevsky, C. M. Cuerrier and A. E. Pelling, PLoS One, 2016, 11(6), e0157894.

47. J. Gershlak, S. Hernandez, G. Fontana, L. Perreault, K. Hansen, S. Larson … G. Gaudette, Biomaterials, 2017, 125, 13–22.

48. S. Thorsteinsdóttir, M. Deries, A. S. Cachaço and F. Bajanca, Dev. Biol., 2011, 354(2), 191–207.

49. K. Novotna, P. Havelka, T. Sopuch, K. Kolarova, V. Vosmanska, V. Lisa … L. Bacakova, Cellulose, 2013, 20(5), 2263–2278.

50. R. J. Hickey and A. E. Pelling, Front. Bioeng. Biotechnol., 2019, 7, 45.

51. J. C. Courtenay, C. Deneke, E. M. Lanzoni, C. A. Costa, Y. Bae, J. L. Scott and R. I. Sharma, Cellulose, 2018, 25(2), 925–940.

52. J. C. Courtenay, M. A. Johns, F. Galembeck, C. Deneke, E. M. Lanzoni, C. A. Costa … R. I. Sharma, Cellulose, 2017, 24(1), 253–267.

53. E. Scarpella and A. Meijer, New Phytol., 2004, 164(2), 209–242.

54. G. N. Karam, Ann. Bot. (Oxford, U.K.), 2005, 95(7), 1179–1186.

55. A. A. Myburg, S. Lev‐Yadun and R. R. Sederoff, eLS, 2013, DOI: 10.1002/9780470015902.a0001302.pub2.

56. J. A. Zallen, Cell, 2007, 129(6), 1051–1063.

57. M. D. Murtey and P. Ramasamy, in Modern Electron Microscopy in Physical and Life Sciences, ed. M. Janecek and R. Kral, IntechOpen, London, U.K., 2016, 173–175.

58. W. Roman and E. R. Gomes, Sem. Cell Dev. Biol., 2018, 82, 51–56.

59. M. Schuetz, R. Smith and B. Ellis, J. Exp. Bot., 2013, 64(1), 11–31.

60. J. D. Szustakowski, J. H. Lee, C. A. Marrese, P. A. Kosinski, N. R. Nirmala and D. M. Kemp, Genomics, 2006, 87(1), 129–138.

61. L. T. Denes, L. A. Riley, J. R. Mijares, J. D. Arboleda, K. McKee, K. A. Esser and E. T. Wang, Skeletal Muscle, 2019, 9(1), 17.

62. V. Hosseini, S. Ahadian, S. Ostrovidov, S. Chen, M. Ramalingam, A. Khademhosseini… H. Kaji, Tissue Eng., Part A, 2012, 18(23–24), 2453–2465.

63. M. Griffin, S. Sen, H. Sweeney and D. Discher, J. Cell Sci., 2004, 117(24), 5855–5863.

64. G. De La Haba, H. M. Kamali and D. M. Tiede, Proc. Natl. Acad. Sci. U. S. A., 1975, 72(7), 2729–2732.

65. K. E. Alley and F. F. Omerza, F. F. Cells Tissues Organs, 1999, 164(1), 46–58.

66. M. J. Perry, J. Tait, J. Hu, S. C. White and S. Medler, J. Exp. Biol., 2009, 212(Pt 5), 673–683.

67. D. Pette and R. S. Staron, Int. Rev. Cytol., 1997, 170, 143–223.

68. K. R. Dalrymple, T. I. Prigozy and C. F. Shuler, C. F. Differentiation; Research in Biological Diversity, 2000, 66(4–5), 218–226.

69. E. Soliman, F. Bianchi, J. N. Sleigh, J. H. George, M. Z. Cader, Z. Cui and H. Ye, Biotechnol. Lett., 2018, 40(3), 601–607.

70. A. D. Schoenenberger, J. Foolen, P. Moor, U. Silvan and J. G. Snedeker, Acta Biomater., 2018, 71, 306–317.

71. N. F. Huang, R. J. Lee, S. Li, S. Am. J. Transl. Res., 2010, 2(1), 43.

72. C. P. Pennisi, C. G. Olesen, M. de Zee, J. Rasmussen and V. Zachar, Tissue Eng., Part A, 2011, 17(19–20), 2543–2550.

73. J. D. Glawe, J. B. Hill, D. K. Mills and M. J. McShane, J. Biomed. Mater. Res., Part A, 2005, 75(1), 106–114.

74. L. Hindi, J. D. McMillan, D. Afroze, S. M. Hindi and A. Kumar, Bio-Protoc., 2017, 7(9).

75. K. Tanaka, K. Sato, T. Yoshida, T. Fukuda, K. Hanamura, N. Kojima … H. Watanabe, Muscle Nerve, 2011, 44(6), 968–977.

76. R. H. Farahi, A. M. Charrier, A. Tolbert, A. L. Lereu, A. Ragauskas, B. H. Davison and A. Passian, Sci. Rep., 2017, 7(1), 152.

77. D. G. Hepworth and J. F. V. Vincent, Ann. Bot. (Oxford, U.K.), 1998, 81(6), 751–759.

78. D. R. Lee, J. Exp. Bot., 1981, 32(126), 251–260.

79. A. J. Engler, M. A. Griffin, S. Sen, C. G. Bönnemann, H. L. Sweeney and D. E. Discher, J. Cell Biol., 2004, 166(6), 877–887.

80. M. Levy-Mishali, J. Zoldan and S. Levenberg, Tissue Eng., Part A, 2009, 15(4), 935–944.

81. R. G. Wells, Hepatology, 2008, 47(4), 1394–1400.

82. A. Lourenço, J. Rencoret, C. Chemetova, J. Gominho, A. Gutiérrez, J. C. del Río and H. Pereira, Front. Recent Dev. Plant Sci., 2016, 7, 1612.

83. B. J. Papenburg, E. D. Rodrigues, M. Wessling and D. Stamatialis, Soft Matter, 2010, 6(18), 4377–4388.

84. B. Cadot, V. Gache, V and E. R. Gomes, Nucleus, 2015, 6(5), 373–381.

85. J. Gingras, R. M. Rioux, D. Cuvelier, N. A. Geisse, J. W. Lichtman, G. M. Whitesides and J. R. Sanes, Biophys. J., 2009, 97(10), 2771–2779.

86. M. P. Olivieri, K. H. Kittle, K. S. Tweden and R. E. Loomis, Biomaterials, 1992, 13(4), 201–208.

