## Supporting Info for "Decellularized Plant-Based Scaffolds for Guided Alignment of Myoblast Cells"

Andrew E. Pelling

University of Ottawa

STEM complex

150 Louis-Pasteur Private

Ottawa, ON K1N 6N5

Canada

Web: <http://www.pellinglab.net>

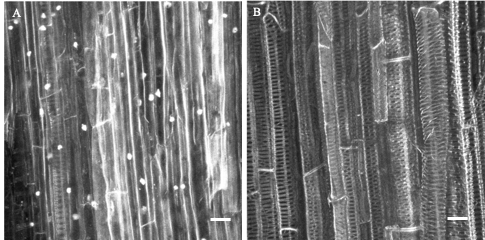

Figure S1. Hoescht 33342 staining of (A) native and (B) decellularized vascular bundle. Nuclei correspond to companion cells of phloem. Scale bar = 25μm.

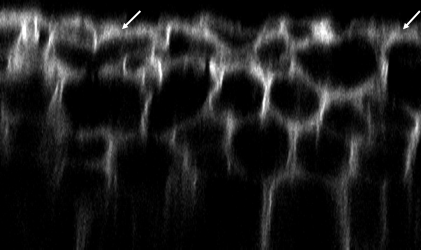

Figure S2. Orthogonal view of phloem microgrooves stained with calcoflour. The image was taken using a multiphoton microscope. Arrows: smooth areas. 184 x 109 μm orthogonal view

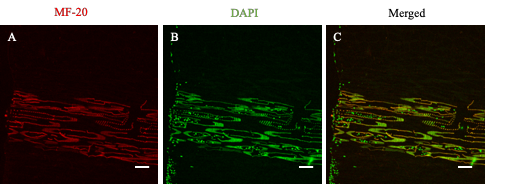

Figure S3. Immunohistochemical analysis of MYHC expression in C2C12 cultured on intact vascular bundles of cross section scaffolds. (A)Myotubes were labeled with MF-20 followed by goat anti-mouse-568 (Red); (B) nuclei were labeled with DAPI (Green). Ribbed structures correspond to fluorescence emission from vascular bundle. Scale bar = 100μm.
